# Ventromedial prefrontal cortex supports prototype representations in healthy older adults

**DOI:** 10.1101/2025.09.24.678354

**Authors:** Caitlin R. Bowman, Dagmar Zeithamova

**Author notes:** Correspondence: Dagmar Zeithamova, PhD.

## Abstract

The ability to learn broad concepts from individual instances is relevant throughout our lifespans as new concepts enter the world, and we seek to acquire new skills and hobbies that can enrich our lives. While older age has been associated with declines in the ability to remember individual instances, less is known about how these declines impact concept learning and generalization or the neural systems that older adults recruit to support abstraction. In the present study, we used prototype-based category learning as a domain to test age differences in concept learning. Young and older adults completed a category-learning task while undergoing fMRI. We fit formal prototype and exemplar models to behavioral and brain data to index concept learning based on abstraction versus memory for individual category members. We found that the fit of both models to behavior was poorer in older adults, but older adults were more likely than young adults to be best fit by the prototype model and less likely to be best fit by the exemplar model. While only young adults showed significant prototype-tracking in the hippocampus, both young and older adults recruited the ventromedial prefrontal cortex (VMPFC) to support prototype-based generalization. Although evidence for age differences in prototype representations emerged in a whole-brain analysis, evidence for age differences were weak in the VMPFC and hippocampus. Thus, engagement of the VMPFC prototype-learning system may help maintain concept generalization in older adults.

## 1. Introduction

It is well established that decline in the ability to remember details of individual experiences is part of normal aging (Alghamdi and Rugg, 2020; Hashtroudi et al., 1989; Prull et al., 2006; Stark et al., 2010). Yet, remembering specific experiences is not the only way we use memory; we also use memory to generalize across experiences to learn new concepts (Morton and Preston, 2021; Schapiro et al., 2017; Zeithamova and Bowman, 2020). Yet, how older age affects memory generalization is not well understood. On the one hand, some memory theories posit that the same memory representations underlie the ability to remember specific episodes and the ability to generalize (Hintzman, 1984; Kumaran and McClelland, 2012; Nosofsky, 1988), in which case we would expect declines in memory specificity to also have negative consequences for older adults’ generalization abilities. On the other hand, there are a constellation of memory abilities that are relatively intact in older age that could, in theory, support generalization. Older adults tend to encode common features across experiences and extract their meaning, or ‘gist’ (Brainerd and Reyna, 2015, 2002) rather than the details that distinguish between similar experiences (Bowman et al., 2019; Bowman and Dennis, 2015; Gallo et al., 2006; Koutstaal and Schacter, 1997). Partial overlap with prior experience is more likely to trigger retrieval (i.e., pattern completion) in older compared to young adults (Vieweg et al., 2015; Wilson et al., 2006; Wynn et al., 2021). These findings point to ways that older adults’ memory abilities may be well suited to forming memory representations that integrate across experiences, which could then support concept learning and generalization.

Because categorization typically involves both learning individual category members and generalization to new examples, it is well suited to addressing how older age impacts the ability to form new conceptual knowledge. Aging studies of categorization have generally shown age deficits in categorization performance, with the degree of the deficit differing by category structure and task (for review see Bowman et al., 2023). Studies have shown small to no age deficit in learning typical category members, but larger age deficits for atypical or exception items (Bowman et al., 2022; Davis et al., 2012; Valdez et al., in press). Prior work has also shown that age differences in categorization arise primarily in learning, with little to no evidence of a generalization-specific deficit (Bowman et al., 2021) and sometimes even age-matched generalization performance despite deficits in learning performance (Bowman et al., 2022; Valdez et al., in press). Thus, there is some evidence that older adults learn individual category members less well than young adults, but that this deficit does not always have a major impact on their ability to generalize.

Further, categorization is a useful domain for testing representations underlying memory judgments because there are longstanding models of categorization that differentially emphasize memory specificity versus integration and abstraction. The exemplar model posits that categories are represented by individual category members stored in memory and that generalization involves joint retrieval of these individual category members to determine the similarity between the to-be-categorized example and the stored examples (Medin and Schaffer, 1978; Nosofsky, 1991). The prototype model, in contrast, posits that categories are represented by prototypes – an idealized category representation that is abstracted from the individual examples (Minda and Smith, 2011; Posner and Keele, 1968). Prototype-based generalization involves determining the similarity between a to-be-categorized example and relevant category prototypes. Based on older adults’ relatively intact gist-based memory and declines in memory specificity, one might expect older adults to be more likely than young adults to rely on prototype representations. Yet, the use of formal exemplar versus prototype modeling in aging studies has been limited, and the findings have been mixed. Formal model fits have sometimes pointed to older adults being less likely than young adults to rely on an exemplar strategy and/or more likely to rely on prototypes (Bowman et al., 2022; Mata et al., 2012). Other findings showed similar strategies across age groups (Mata et al., 2012; Schenk et al., 2016; Valdez et al., in press). Thus, further evidence is needed to understand how older age affects prototype- and exemplar-based categorization.

Another advantage of the categorization domain is that latent variables from the prototype and exemplar models can be fit to fMRI data, allowing researchers to test whether the brain tracks the information needed to make prototype- or exemplar-based categorization judgments. In young adults, prototype correlates have been identified in the ventromedial prefrontal cortex (VMPFC) and the anterior hippocampus (Bowman et al., 2020; Bowman and Zeithamova, 2018; Liu et al., 2025). These findings are in line with the VMFPC and hippocampus playing a role in memory integration across multiple memory domains (Shohamy and Wagner, 2008; van Kesteren et al., 2010; Zeithamova et al., 2012). In contrast, exemplar correlates have been identified in occipital and parietal regions (Blank and Bayer, 2022; Bowman et al., 2020; Mack et al., 2013), inferior lateral prefrontal cortex (Bowman et al., 2020; Mack et al., 2013), and the hippocampus (Blank and Bayer, 2022). Notably, while some studies have shown only prototype (Bowman and Zeithamova, 2018) or only exemplar correlates (Mack et al., 2013) in the brain, other work has shown both prototype and exemplar correlates in the brain in the same task (Bowman et al., 2020) and even within the same individuals (Blank and Bayer, 2022). Thus, brain imaging can provide novel information about the kinds of representations available to make generalization judgments, even when one representational strategy is most prominent in behavior. Yet, prototype and exemplar models have not been fit to fMRI data in older adults, leaving the neural underpinnings of their categorization judgments unclear.

In the present study, we aimed to test the extent to which older adults recruit the same regions as young adults to support prototype-based categorization (the VMPFC and anterior hippocampus), and the extent of any age differences in prototype correlates in those regions. To do so, young (aged 18-30) and older adults (aged 60+) learned and were tested on a prototype-based category structure using novel cartoon animals that have 10 binary features (Figure 1A). In this structure, one cartoon serves as the prototype of category A and has the most common version of each feature from the category members. The prototype of category B is the stimulus that shares no features with the category A prototype. Items that share more features with the category A prototype than the category B prototype are considered members of category A and vice versa. With this structure, we can determine the number of shared features between any pair of stimuli, allowing for computation of prototype- and exemplar-based similarity as needed by the formalizations of the prototype and exemplar models. Participants completed a feedback-based category training task outside the scanner (Figure 1B), then underwent fMRI scanning during a categorization test that included both training items and new generalization items but no feedback (Figure 1C). We fit formal prototype and exemplar models to the behavioral categorization test responses and to brain activation levels measured with fMRI. We tested the extent of age differences in category learning and generalization performance, the extent to which behavioral responses were best explained by the prototype versus exemplar model, and the extent to which signals in the anterior hippocampus and VMPFC correlated with predictions from the prototype model.

**Figure 1.**
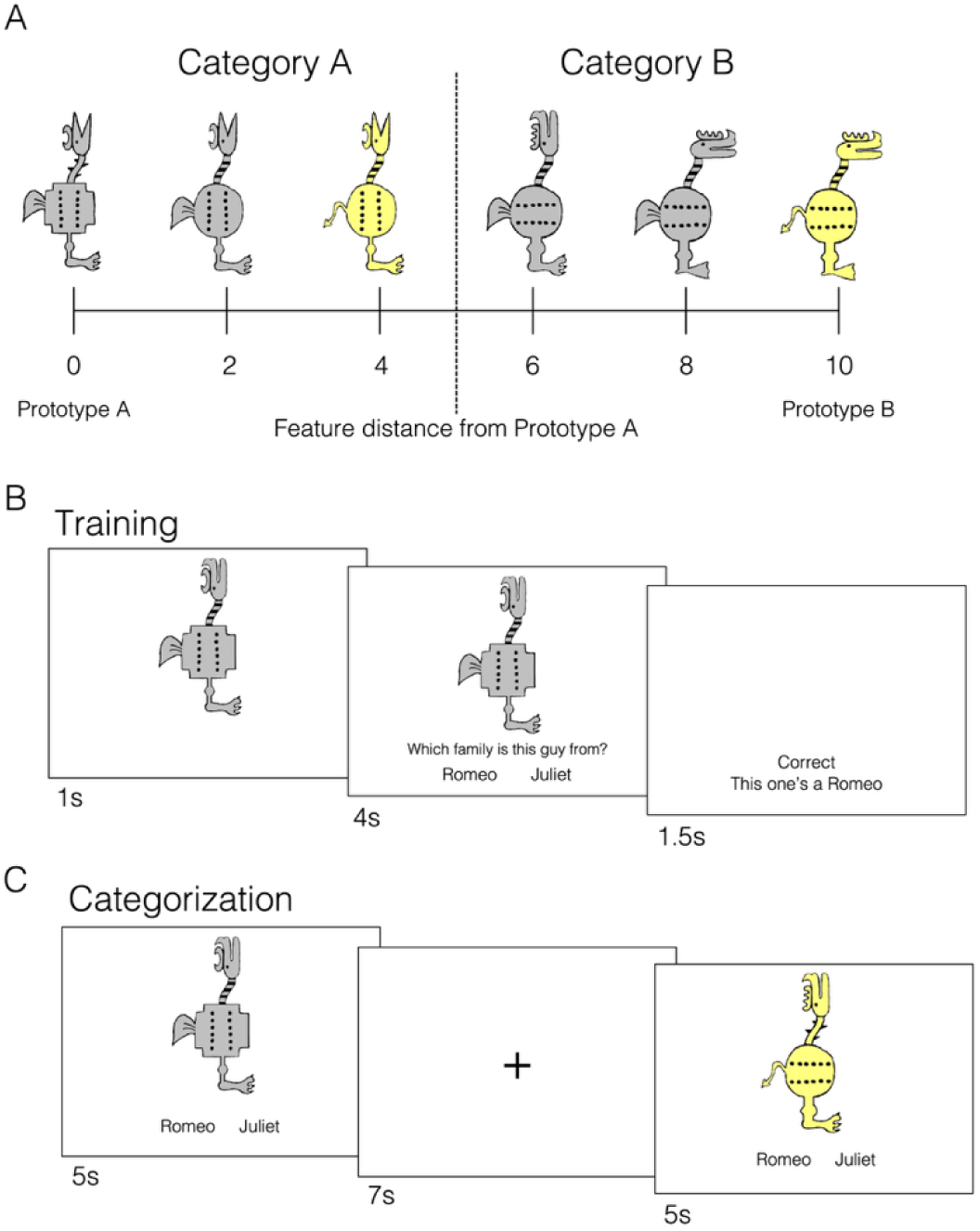
Category structure and task procedure. A. The categories were designed around category prototypes. The prototypes had the typical version of each feature from the training items of their category. Items that shared more features with the category A prototype were members of category A (“Romeo”), and vice versa for category B (“Juliet”). B. Outside the scanner, participants completed feedback-based training with a set of eight distance 2 items and six distance 4 items. Immediately after training, participants also completed a brief old/new recognition test outside the scanner (not depicted). C. In the scanner, after structural scans and a rest scan, participants completed four runs of a categorization task. Participants categorized training items and new items without feedback. *Note*: s = seconds.

## 2. Method

### 2.1. Data and Code Availability

The de-identified behavioral and fMRI data and the analytic code are publicly available through OpenNeuro (Reviewer Link). The study was not pre-registered.

### 2.2. Participants

Forty-two young adults (recruitment range = 18-30 years) and forty older adults (recruitment range = 60+ years) were recruited from the University of Oregon and surrounding community and participated for financial compensation. This sample size was determined based on the ability to detect a small-medium effect size (d = 0.4) for prototype correlates in the anterior hippocampus and VMPFC in each age group, based on the strength of these effects in young adults in prior studies (Bowman et al., 2020; Bowman and Zeithamova, 2018) (one-tailed, one-sample t-test, alpha = .05, Power = 80%). After exclusions (see below), our samples were powered to detect effects of d > .48 within the older group, d > .42 within the younger group, and two-tailed age differences of d > .7. All experimental procedures were approved by Research Compliance Services at the University of Oregon, and all participants provided written informed consent. Participants were screened to be right-handed, English-speaking, free of self-reported neurological conditions, and not taking medications known to affect brain function (see Supplementary materials for full screening protocol). Older adults were also screened for signs of cognitive impairment using the Mini Mental State Exam (score > 24 for inclusion). All participants scored above this criterion (M = 28.7, SD = 1.3, range 26-30 for those included in analyses). One young adult and two older adults were excluded from analyses due to technical issues with the scan session that prevented data collection. One challenge with using prototype and exemplar-based regressors in fMRI analysis is that they are based on participants’ responses rather than pre-defined conditions. In some cases, the two models’ predictions end up highly correlated (Bowman et al., 2020). Five young adults and nine older adults were excluded for having fMRI model regressors that were highly correlated in three or more runs (r > .9). Following exclusions, 36 young adults (mean age = 20.5 years, range = 18-27, 24 females) and 28 older adults (mean age = 70.1 years, range = 62-86 years; 15 females) were included in all analyses. An additional four young adults and ten older adults had excessively correlated regressors in one or two runs. We retained those participants but excluded those individual runs from analysis.

### 2.3. Materials

In generating stimulus sets, pairs of young and older adults were yoked so that both the stimuli themselves and their presentation order were the same, minimizing the potential for item or order effects to drive age differences. Stimuli were novel cartoon animals with 10 binary dimensions (https://osf.io/8bph2/) (Bowman and Zeithamova, 2020; Bozoki et al., 2006). The two versions of each feature are visible across the example category A and category B prototypes in Figure 1A. For each participant, one stimulus was chosen to be the prototype of category A from a list of four possible prototypes, with prototypes counterbalanced across participants. The stimulus sets were designed such that the category A prototype would have all the most typical versions of each feature based on the category A members. All stimuli were coded in reference to the category A prototype. The stimulus that shared no features with the category A prototype was the category B prototype, which similarly had all the most typical feature values based on the category B members. Category A stimuli were those that shared more features with the category A prototype than the category B prototype. Category B stimuli were those that shared more features with the category B prototype than the category A prototype. Stimuli sharing an equal number of features with each prototype were not used in the study.

#### 2.3.1. Training set

The training set included items that were near the category center (differing by two of ten features from their category prototype, ‘distance 2’ items) and items that were near the category boundary (differing by four of ten features from their category prototype, ‘distance 4’ items). This feature of the training sets allowed us to test whether there were larger age differences in learning category typical (distance 2) versus category atypical (distance 4) items, as we have found previously (Bowman et al., 2022; Valdez et al., in press). We also chose a structure that had previously shown balanced prototype versus exemplar representations in young adults (Bowman and Zeithamova, 2020), allowing room for age-related increases or decreases in prototype representations. The prototypes themselves were not included in the training set. Training sets included seven items per category (14 total items across categories): four distance 2 items and three distance 4 items per category (see Table 1 for the training set structure). These numbers of training items at each distance were selected so that we could constrain the set with each feature equally predictive of category membership so that participants would need to holistically evaluate each stimulus rather than learning to focus on specific features that were most diagnostic. The overall structure of the training set was the same across participants, but the exact stimuli depended on the prototypes selected.

**Table 1.** Category training set structure.

| Stimulus | Dimension values |  |  |  |  |  |  |  |  |  |
| --- | --- | --- | --- | --- | --- | --- | --- | --- | --- | --- |
|  | 1 | 2 | 3 | 4 | 5 | 6 | 7 | 8 | 9 | 10 |
| Prototype A | 1 | 1 | 1 | 1 | 1 | 1 | 1 | 1 | 1 | 1 |
| A1 | 0 | 0 | 1 | 1 | 1 | 1 | 1 | 1 | 1 | 1 |
| A2 | 1 | 1 | 0 | 0 | 1 | 1 | 1 | 1 | 1 | 1 |
| A3 | 1 | 1 | 1 | 1 | 0 | 0 | 1 | 1 | 1 | 1 |
| A4 | 1 | 1 | 1 | 1 | 1 | 1 | 0 | 0 | 1 | 1 |
| A5 | 0 | 1 | 1 | 0 | 1 | 1 | 1 | 0 | 1 | 0 |
| A6 | 1 | 0 | 1 | 1 | 0 | 1 | 0 | 1 | 0 | 1 |
| A7 | 1 | 1 | 0 | 1 | 1 | 0 | 1 | 1 | 0 | 0 |
| Prototype B | 0 | 0 | 0 | 0 | 0 | 0 | 0 | 0 | 0 | 0 |
| B1 | 1 | 1 | 0 | 0 | 0 | 0 | 0 | 0 | 0 | 0 |
| B2 | 0 | 0 | 1 | 1 | 0 | 0 | 0 | 0 | 0 | 0 |
| B3 | 0 | 0 | 0 | 0 | 1 | 1 | 0 | 0 | 0 | 0 |
| B4 | 0 | 0 | 0 | 0 | 0 | 0 | 1 | 1 | 0 | 0 |
| B5 | 1 | 0 | 0 | 1 | 0 | 0 | 0 | 1 | 0 | 1 |
| B6 | 0 | 1 | 0 | 0 | 1 | 0 | 1 | 0 | 1 | 0 |
| B7 | 0 | 0 | 1 | 0 | 0 | 1 | 0 | 0 | 1 | 1 |

*Note:* Prototypes were not included in the training set but are depicted here to illustrate the relationship between training items and the prototypes. A1-A7 represent category A training items. B1-B7 represent category B training items. Items 1-4 of each category were distance 2, and items 5-7 were distance 4.

#### 2.3.2. Categorization set

Sixty-four unique items were included in the categorization test set. The test included the 14 (old) training items, each presented twice. We presented them twice because they are limited in number and critical items for fitting the exemplar model: the exemplar model predicts that categorization decisions are based on similarity to the training items, so the training items themselves should typically be categorized especially well. A total of 50 unique new items were included in the test set as a measure of category generalization. New items included the two prototypes, each presented twice because they are limited in number and critical for fitting the prototype model (Bowman and Zeithamova, 2018; Kéri et al., 2001; Smith et al., 2008). An additional 48 new items were six new stimuli from each distance from the category A prototype (1-4, 6-9) each presented once. These new items were selected randomly for each yoked participant pair from the items not included in the training set.

### 2.4. Procedure

#### 2.4.1. Overview

Participants completed the training phase and a brief old/new recognition test (not reported here) outside the scanner. They were then given information about the MRI environment, instructions for the categorization test, and were setup in the scanner. The scan began with a high resolution T1-weighted anatomical scan, followed by a resting state scan (not reported here). The categorization test followed the rest scan and was split across four runs. The scan ended with a T2-weighted anatomical image. After the scan, participants were given questionnaires about their strategies during the task, debriefed, and given their compensation.

#### 2.4.2. Training

In each trial of the training task (Figure 1B), an individual training item was presented, and participants were instructed to decide which of two families (Romeo’s family or Juliet’s family) they thought the animal belonged to. The stimulus appeared on the screen alone for 1 second before the response options appeared, giving participants time to evaluate the stimulus without needing to respond. One second into the trial, the response options appeared alongside the stimulus. The stimulus remained on the screen for an additional 4 seconds (5 seconds total) regardless of when the participant responded so that all participants would have equal exposure to the stimulus. Once the participant responded, corrective feedback (“CORRECT” or “WRONG”) appeared immediately alongside the item’s correct family membership (“It was Romeo’s family” or “It was Juliet’s family”). If the response was made while the stimulus was on the screen, then the feedback replaced the response options and appeared alongside the stimulus. This procedure ensured that feedback was immediate, and that participants had equal exposure to the stimuli regardless of how quickly they responded. The feedback then stayed on the screen for an additional 1.5 seconds after the stimulus disappeared. If the participant did not respond while the stimulus was on the screen, they were prompted to respond immediately (“Please respond now”) and given an additional 1 second to respond. If they responded within the 1 second, the corrective feedback was presented for 1.5 seconds. If they did not respond in time, the feedback “Respond faster” was presented with the item’s family membership for 1.5 seconds. Each trial was followed by a 1-second fixation cross presented.

There were eight blocks of training separated by self-paced breaks. Each block contained four repetitions of each of the 14 training items, totaling 56 trials per block and 448 total training trials. Within each block, the order of stimuli was randomized with the constraint that all training items were presented before any were presented again and so that no more than three items from the same category were presented consecutively.

#### 2.4.3. Categorization

The categorization test took place in the scanner following an anatomy scan and a rest scan. On each trial, an individual item was presented, and participants were asked to decide which family they thought it belonged to. The response options appeared immediately alongside the stimulus, and participants had 5 seconds to make their response while the stimulus was still on the screen. There was no feedback during the test phase. Each trial was followed by a 7-second fixation cross (i.e., slow-event related design (Dale, 1999; Liu, 2012; Zeithamova et al., 2017)) in line with our previous work in young adults (Bowman et al., 2020; Bowman and Zeithamova, 2018). The 80 categorization trials were split evenly across four scan runs. Old items, prototypes, and new items at each distance were approximately evenly distributed across the four runs. The presentation order was randomized with the constraints that no more than two old items and no more than three items from the same category were presented consecutively.

### 2.5. MRI data acquisition

MRI data were acquired on a 3T Siemens MAGNETOM Skyra scanner at the University of Oregon Lewis Center for Neuroimaging. A 32-channel head coil was used. We minimized head motion with foam padding, monitored head motion using FIRMM online motion tracking software (Turing Medical, https://turingmedical.com/firmm/), and gave participants feedback about how well they held still in between runs.

The scanning session started with a localizer scan followed by a standard, high-resolution T1-weighted MPRAGE anatomical image (TR = 2500 ms; TE = 3.43 ms; TI = 1100 ms; flip angle = 7°; matrix size = 256 x 256; 176 contiguous slices; FOV = 256 mm; slice thickness = 1 mm; voxel size = 1.0 x 1.0 x 1.0 mm; GRAPPA factor = 2). Five runs of functional data collection followed using a multiband gradient echo pulse sequence (TR = 2000 ms; TE = 25 ms; flip angle = 90°; matrix size = 104 x 104; 72 contiguous slices oriented 15° off the anterior-posterior commissure line to reduce prefrontal signal dropout; FOV = 208 mm; voxel size = 2.0 x 2.0 x 2.0 mm; GRAPPA factor = 2; multiband acceleration factor = 3). The first functional run was a resting state scan (180 volumes lasting 6 minutes) followed by four categorization runs (122 volumes and lasting 4 minutes and 4 seconds each). The scan session ended with a T2-weighted coronal anatomical image (TR =13,520 ms; TE = 88 ms; flip angle = 150°; matrix size = 512 x 512; 65 contiguous slices manually oriented perpendicularly to the hippocampal long axis; interleaved acquisition; FOV = 220 mm; voxel size = 0.4 x 0.4 x 2 mm; GRAPPA factor = 2).

### 2.6. MRI data pre-processing

#### 2.6.1. Regions of interest

T1-weighted anatomical scans were processed using Freesurfer version 6.0.0 to generate cortical and subcortical regions of interest (ROIs) in each participant’s native space. Because we did not have predictions about laterality effects, we used ROIs that averaged across the right and left hemispheres. We focused on the ventromedial prefrontal cortex (medial orbitofrontal Freesurfer label) and anterior hippocampus based on our prior work in young adults showing prototype correlates in these regions (Bowman et al., 2020; Bowman and Zeithamova, 2018). We also included the posterior hippocampus to determine whether the strength of any prototype effects differed across the hippocampal long axis. The anterior versus posterior hippocampus was defined based on the middle slice, with the middle slice going to the posterior hippocampus when there was an odd number of slices. After anatomical definition in structural space, all ROI masks were registered to functional space using Advanced Normalization Tools (ANTs; https://stnava.github.io/ANTs/).

#### 2.6.2. fMRI pre-processing

Raw dicom images were converted to Nifti format using the dcm2nii function from MRIcron (https://www.nitrc.org/projects/mricron). Skull tissue was removed from the functional images using FSL’s brain extraction tool (BET) (www.fmrib.ox.ac.uk/fsl). Within-run motion correction was computed and applied using FSL’s MCFLIRT, realigning all volumes to the middle volume. We then used ANTs to align volumes across runs with the first volume of the first functional run (rest) serving as the reference and the first volume of all other runs used to compute the transformation to the reference and then applied to all volumes in each run. We excluded individual runs with any framewise displacement (FD) value > 1.5 mm to ensure all movements were smaller than our voxel size (2 mm^3^). All participants had at least two runs of data included. To determine whether there were age differences in motion estimates after motion exclusions, we conducted a linear mixed effects model with the maximum FD value for each participant in each run as the dependent variable. Unsurprisingly, there was a significant effect of age group (ß = .29, *t* = 8.91, *p* < .001). Nonetheless, the mean of the maximum FD values for older adults was still quite good (M = .64 mm, SD = .25 mm) although higher than the excellent mean value for young adults (M = .35 mm, SD = .13 mm). Further, the proportion of FD values that were < .5 mm was extremely high in both age groups (young M = 99.9%, SD = .5%; older M = 96.8%, SD = 6%). We input the resulting motion corrected images into FSL’s fMRI Expert Analysis Tool (FEAT) for high pass temporal filtering (100 s) and spatial smoothing (4 mm FWHM gaussian kernel).

In addition to ROI analyses conducted in native space, we also conducted exploratory whole-brain analyses. For whole-brain analyses, we registered each participant’s brain to the MNI template (MNI152_T1_2mm_brain) using FSL’s FMRIB’s Linear Image Registration Tool (FLIRT). Functional images were first registered to the high-resolution structural image using a linear transformation, then registered to the MNI template using linear registration with 12 degrees of freedom.

### 2.7. Statistical analyses

#### 2.7.1. Behavioral statistical models

Our overall approach for accuracy analyses was to compute Age Group x Distance mixed factors ANOVAs on the data from the training and categorization phases. Where the sphericity assumption was violated, we used a Greenhouse-Geisser correction (denoted with ‘GG’).

To evaluate participants’ categorization strategies, we fit the exemplar and prototype models to the behavioral categorization responses using the same approach as our prior work (Bowman et al., 2022, 2020; Bowman and Zeithamova, 2023, 2020, 2018), which is described in detail in Appendix A. Briefly, the exemplar model assumes that an item is categorized based on its similarity to members of relevant categories stored in memory. Thus, we compute the similarity of each categorization test item to the training items from each category, taking into account the fact that participants may not attend equally to all stimulus features and that there may be differences across participants in the relationship between the number of differing features and subjective similarity. We then sum the similarity values across all members of each category to generate a single value per category that represents the subjective similarity of the to-be-categorized item to the exemplar-based category A and category B representations. Prototype-based similarity is the same except that the similarity is in reference to the prototype of each category rather than the individual training items. For both models, the probability of categorizing an item as a member of a given category is the similarity to that category divided by the summed similarity across both categories. We generated these probabilities for each model and for each trial of the categorization test, creating model predictions that we then compared to the observed responses using an error metric (negative log likelihood). The model with the lower error had a better fit. We then used a permutation testing approach (Zeithamova et al., 2025) to determine whether each model fit better than one would expect by chance and whether one model fit significantly better than the other.

#### 2.7.2. fMRI statistical models

Prototype and exemplar-based modeling of fMRI data was conducted as in prior work (Bowman et al., 2020; Bowman and Zeithamova, 2018). For each participant, for each of the four categorization runs, we conducted a first-level General Linear Model in FSL’s FEAT that included three regressors of interest: 1) item regressor: all trial onsets with a duration of 5 seconds (i.e., the length of the stimulus presentation) and constant weighting (i.e., weight = 1 for each trial), 2) exemplar regressor: the same as the item regressor but with the trial-by-trial summed similarity (category A similarity + category B similarity from the behavioral model fitting) as estimated by the exemplar model for the weight, and 3) prototype regressor: the same as the item regressor but with the trial-by-trial summed similarity as estimated by the prototype model for the weight. Using the prototype and exemplar summed similarity as weights served to parametrically modulate the predicted BOLD response such that a larger amplitude BOLD response is predicted on trials with higher exemplar- or prototype-based similarity. These onsets were then convolved with a canonical hemodynamic response function (gamma function with a phase of 0 s, an SD of 3 s, and a mean lag time of 6 s) and used as predictors of the observed BOLD response. We also included translational and rotational motion as regressors of no interest.

For ROI analyses, we extracted the mean prototype and exemplar parameter estimates across each ROI from each run. For each model, we then computed the mean parameter estimate across runs, dividing by the standard deviation across runs. Normalizing the parameter estimates by their variability can de-weight values associated with higher uncertainty, similar to how lower-level estimates are used in group analyses in FSL (Smith et al., 2004). The resulting prototype and exemplar effects from each participant were then submitted to group-level analyses. We first conducted one-sample t-tests comparing the prototype effects in the VMPFC and anterior hippocampus in each age group to baseline to determine if young and older adults engaged these regions to support prototype-tracking during generalization. Because of our directional prediction, we used one-tailed tests but also included a correction for the four tests conducted (one-tailed corrected alpha = .0125). For these key comparisons of interest, we also include Bayes Factors to indicate the strength of the evidence in favor of a prototype effect in each region and each age group. We followed up these one-sample tests with two ANOVAs – one for the VMPFC and one for the anterior hippocampus. Both ANOVAs included age group (young, older) and model predictor (prototype, exemplar) as factors. Because prior work in young adults has found prototype correlates to be stronger in the anterior than posterior hippocampus (Bowman et al., 2020; Bowman and Zeithamova, 2018), we also included a hippocampal subregion factor (anterior, posterior) in the ANOVA for the hippocampus. Follow-up t-tests were conducted for significant effects, using two-tailed tests. Given the importance of age effects to our research aims, we computed Bayes Factors to quantify the evidence for or against the presence of age differences in each ROI regardless of the significance of age effects in the ANOVA.

For whole-brain analyses, parameter estimates were averaged across runs within individual participants using a fixed-effects analysis. Group-level contrasts were computed using mixed-effects FLAME 1 and 2 in FSL. We compared the prototype and exemplar regressors to the implicit baseline across the entire sample, within the young adult sample alone, and within the older adult sample alone. We also computed independent samples t-test contrasts comparing young and older adults for prototype and exemplar correlates separately. Whole-brain maps used a voxelwise threshold of *z* > 3.1 and were then cluster corrected at *p* < .05 with the false discovery rate method in FSL.

## 3. Results

### 3.1. Behavioral performance

#### 3.1.1. Training

Mean accuracies for each training block separated by training item distance and age group are presented in Figure 2A. To test for age differences in category learning, we computed an age group (young vs. older) x item distance (2 vs. 4 features different from prototype) mixed factors ANOVA on accuracy in the final training block. There was a significant main effect of age group, *F*(1,62) = 26.37, *p* < .001, *η_p_*^2^ = .30, with young adults reaching higher overall performance than older adults (YA M = .86, SD = .12; OA M = .71, SD = .11). There was also a significant main effect of item distance, *F*(1,62) = 113.07, *p* < .001, *η_p_*^2^ = .65, with better learning of distance 2 items (M = .87, SD = .08) compared to distance 4 items (M = .70, SE = .16). Finally, there was also a significant age group x item distance interaction, *F*(1,62) = 8.90, *p* = .004, *η_p_*^2^ = .13. While young adults significantly outperformed older adults for both item types, that effect was smaller for distance 2 items, *t*(62) = 3.99, *p* < .001, *d* = 1.01, young-old M = .10, compared to distance 4 items, *t*(62) = 4.97, *p* < .001, *d* = 1.25, young-old M = .20. Thus, older adults had more difficulty learning the categories, and that age deficit was particularly prominent for the items furthest from the prototypes.

**Figure 2.**
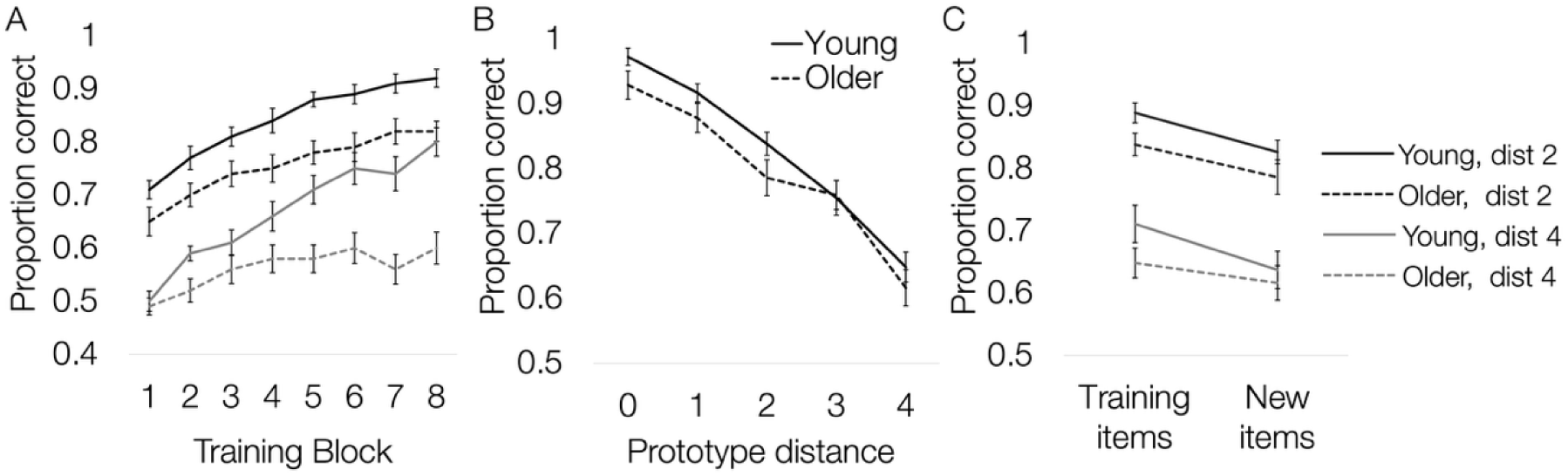
Behavioral accuracy results. A. The proportion of correct responses for each block of training separated by age group and training item distance (dark = distance 2, light = distance 4 as in legend at far right). B. The proportion correct for new items distance 0-4 on the categorization test. C. The proportion correct for training items (old) and new items at distance 2 and 4 on the categorization test. Only distances 2 and 4 are included because all training items came from those distances. For A-C, solid lines represent responses for the young adult group, and dashed lines represent responses for the older adult group. Error bars represent the standard error of the mean across participants. dist = distance.

#### 3.1.2. Categorization test

Mean accuracies for new items presented in the categorization test separated by age group and distance from category prototypes are presented in Figure 2B. While the test included a mix of old (at distance 2 and distance 4) and new (at all distances), we started by focusing on only the new items to test for a prototype gradient in generalization performance. We computed an age group (young, older) x item distance (0-4 features different from the prototype) mixed factors ANOVA on classification accuracy for new test items. Results revealed a significant main effect of item distance, *F*(3.5,217.1) = 80.43, *p* < .001, *η_p_*^2^ = .57, GG, with an accompanying significant linear effect of item distance, *F*(1,62) = 258.29, *p* < .001, *η_p_*^2^ = .06. Generalization performance was worse for items further from category prototypes. There was a non-significant trend toward better generalization in the young group (M = .83, SD = .08) compared to the older group (M = .79, SD = .07), *F*(1,62) = 2.92, *p* = .09, *η_p_*^2^ = .045. Age group and item distance did not significantly interact, *F*(3.5,217.1) = .62, *p* = .64, *η_p_*^2^ = .01, GG.

We also compared accuracies of old and new items as one way to assess the role of exemplar memorization in category decision. We have often found a categorization advantage for old items relative to new items at the same distance from the prototype (Bowman et al., 2022, 2020; Bowman and Zeithamova, 2018), demonstrating some contribution of exemplar memorization even when the prototype model better fits behavior overall. Here we tested whether the old advantage is modulated by age group and item distance (typicality). Mean accuracies separated by age group, item distance (distance 2, distance 4), and item history (training item, new) are presented in Figure 2C. Only distance 2 and 4 items were considered since only those distances had both old and new items. Full results of a mixed factors ANOVA with age group (young, older), item distance (distance 2 or distance 4), and item history (training item, new) are presented in Table 2. As expected, there was a significant main effect of item history, with better classification of training items (M = .77, SD = .10) than new items (M = .72, SD = .10), demonstrating the advantage for old items. With the old items added and only distance 2 and distance 4 items considered, the trend toward an age effect from the generalization became significant, with poorer categorization performance in the older group (M = .72, SD = .08) compared to the young group (M = .77, SD = .08). Finally, there was a main effect of item distance, with better classification of distance 2 items (M = .84, SD = .09) than distance 4 items (M = .65, SD = .13). However, none of the interaction effects reached significance. Thus, while there was categorization advantage for old items over new items indicative of exemplar memory contributing to categorization decisions, this effect was comparable across age groups.

**Table 2.** Age group x Item distance x Item history ANOVA results for categorization accuracy.

| Table 2 |  |  |  |  |
| --- | --- | --- | --- | --- |
| Effect | *df* | *F* value | *p* value | $\eta_p^2$ |
| Age group* | 1,62 | 6.47 | .01 | .10 |
| Item distance* | 1,62 | 115.14 | < .001 | .65 |
| Item history* | 1,62 | 8.59 | .005 | .12 |
| Age group x Item distance | 1,62 | 0.02 | .89 | .00 |
| Age group x Item history | 1,62 | 0.78 | .38 | .01 |
| Item distance x Item history | 1,62 | 0.02 | .90 | .00 |
| Age group x Item distance x Item history | 1,62 | .28 | .60 | .00 |

Note: \* indicates a significant effect at $\alpha = .05$ .

Taken together, behavioral results show a clear age deficit in training accuracy but a less prominent deficit when generalizing. This was especially notable for the category boundary (distance 4) items, where young adults’ training performance ended around 80% but dropped to around 65% during generalization. While older adults’ mean distance 4 training accuracy did not quite reach 60%, generalization accuracy was slightly above 60%, meaning that there was no performance cost for generalization in older adults for items at that distance. We also show a slight performance advantage for classifying training items relative to new generalization items without strong age-related moderation.

### 3.2. Prototype and exemplar model fits

#### 3.2.1. Fits to behavioral categorization responses

The group-level fit values for the prototype and exemplar models are presented in Figure 3A separately for young and older adults. An Age group x Model ANOVA on the fit values revealed a significant main effect of Age group, *F*(1,62) = 11.10, *p* = .001, *η_p_*^2^ = .15, with poorer overall fit in older adults (M = 23.74, SD = 7.7, higher values mean poorer fit) compared to young adults (M = 17.27, SD = 7.7). There was also a significant main effect of model, *F*(1,62) = 9.13, *p* = .004, *η_p_*^2^ = .13, with better fit of the prototype model (M = 19.40, SD = 7.7) compared to the exemplar model (M = 21.61, SD = 8.9). The Age group x Model interaction effect did not reach significance, *F*(1,62) = 1.73, *p* = .19, *η_p_*^2^ = .03. Thus, the prototype model fit both the young and older groups better than the exemplar model, and both models better explained the categorization responses of young adults than those of older adults.

**Figure 3.**
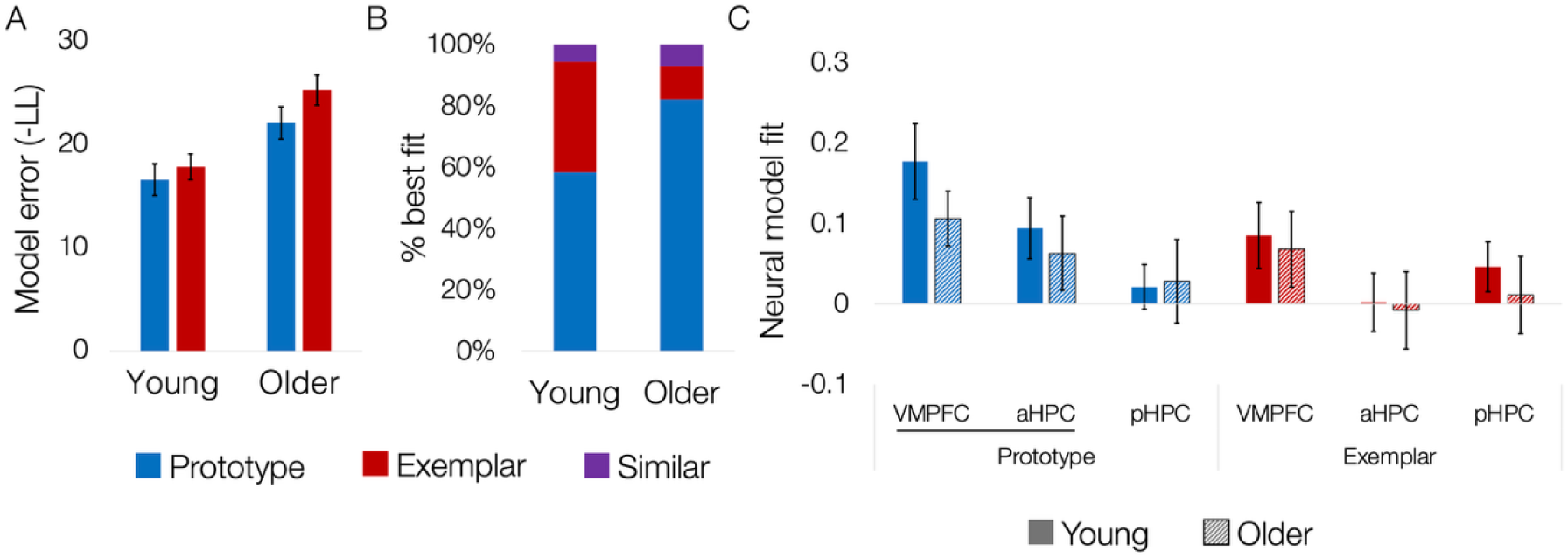
Prototype and exemplar model fits to behavior and neural regions of interest. A. The degree of model error (in terms of negative log likelihood or ‘-LL’) from the fitted prototype and exemplar models separated by age group. B. The percentage of young and older adults whose behavioral responses were best fit by the prototype model, exemplar model, and those who had similar fits for the two models. C. The neural fit of the prototype and exemplar models in the ventromedial prefrontal cortex (VMPFC), anterior hippocampus (aHPC), and posterior hippocampus (pHPC) depicted separately for young and older adults. Prototype effects in the VMPFC and aHPC are underlined as a priori effects of interest. The neural model fit was the mean beta parameter for the model regressor in a given region divided by the standard deviation (i.e., an effect size). In A and C, error bars represent the standard error of the mean across participants.

From these fit values, we determined which model fit best in each individual participant, testing whether a higher proportion of older adults were best fit by the prototype model compared to young adults. Our first step in model selection was to determine whether each model fit better than simply assuming that the participant responded randomly. In the present study, both the prototype and exemplar model fit better than a random model in all participants. Next, we compared the prototype and exemplar models to one another. The proportions of young and older adults best fit by the prototype model, best fit by the exemplar model, and similarly fit by the two models (“similar”) are depicted in Figure 3B. A chi-squared test revealed that the prototype model was the best fitting model in a significantly higher proportion of older adults (82%, n = 23 of 28) compared to young adults (58%, n = 21 of 36), χ^2^ (1) = 4.16, *p* = .04. We also found that the exemplar model was the best fitting model in a significantly higher proportion of young adults (36%, n = 13 of 36) compared to older adults (11%, n = 3 of 28), χ^2^ (1) = 5.42, *p* = .02. Taken together, we found that both the prototype and exemplar model fit more poorly in older adults compared to young adults, but that when the models were compared directly for each participant, older adults were biased toward a prototype strategy whereas young adults showed more mixed prototype and exemplar strategy use.

#### 3.2.2. Neural model fits: regions of interest

Mean prototype and exemplar responses from the anterior and posterior hippocampus and VMPFC are presented separately for young and older adults in Figure 3C. We first tested whether young and older adults recruited the VMPFC and anterior hippocampus to track prototype-based similarity with one-sample prototype > baseline comparisons (one-tailed alpha corrected for four comparisons = .0125). We replicated our prior findings by showing that the VMFPC, *t*(35) = 3.78, *p* < .001, *d* = 0.63, and anterior hippocampus, *t*(35) = 2.50, *p* = .009, *d* = 0.42, tracked prototype predictions in young adults. Bayes Factors indicated very strong evidence in favor of the VMPFC effect (BF_10_ = 100.0) and moderate evidence in favor of an effect in the anterior hippocampus (BF_10_ = 5.3). One-sample t-tests in the older adult group revealed significant prototype tracking in the VMPFC, *t*(27) = 3.12, *p* = .002, *d* = .59, with strong evidence in favor of this effect (BF_10_ = 18.9). Older adults showed a numerically positive but non-significant prototype effect in the anterior hippocampus, *t*(27) = 1.37, *p* = .09, *d* = 0.26, with weak evidence in favor of an effect (BF_10_ = 0.8). Thus, consistent with our predictions, we found strong to very strong evidence of prototype tracking in the VMPFC across age groups. Evidence of prototype tracking was moderate to weak for the anterior hippocampus and did not reach significance in the older group while being in the predicted direction.

We followed up on these prototype effects by testing whether prototype correlates were significantly stronger than exemplar correlates and whether any effects differed across age groups. First, we conducted an Age group (young, older) x Model (prototype, exemplar) mixed effects ANOVA in the VMPFC. The main effect of age was not significant, *F*(1,62) = 1.42, *p* = .24, *η_p_*^2^ = .02, nor was there a significant age x model interaction, *F*(1,62) = .30, *p* = .59, *η_p_*^2^ = .005. Bayes Factors additionally indicated weak evidence in favor of an age difference in either prototype (BF_10_ = 0.5) or exemplar tracking (BF_10_ = 0.3) in the VMPFC. Because there were age differences in motion estimates (small in absolute magnitude but significant), we tested whether spurious motion-related activation in older adults could be masking true age differences in VMPFC prototype correlates by computing a multiple regression with age group and the mean maximum FD across runs predicting VMPFC prototype activation. The age effect remained non-significant when accounting for group differences in motion, *ß* = .01, *SE* = .09, *t* = .12, *p* = .91, and the effect of motion was directionally negative (i.e., the opposite of what could potentially mask age deficits), *ß* = -.28, *SE* = .23, *t* = -1.18, *p* = .24. Thus, we found that prototype effects in the VMPFC did not differ significantly across age groups.

Surprisingly, the main effect of prototype vs. exemplar model was also not significant, *F*(1,62) = 1.75, *p* = .19, *η_p_*^2^ =.03. Given prior findings in young adults showing that the VMPFC tracks prototype information significantly more than exemplars (Bowman et al., 2020; Bowman and Zeithamova, 2018), the lack of a main effect of model was unexpected. We followed up on the lack of a prototype model advantage in the VMPFC by testing whether there was significant exemplar tracking in the VMPFC in either age group. One-sample t-tests revealed exemplar tracking in young adults that passed a conventional alpha = .05 threshold but not a correction for two comparisons (young, older alpha = .025), *t*(35) = 2.08, *p* = .045, *d* = .35. The effect was not significant at either alpha for older adults, *t*(28) = 1.45, *p* = .16, *d* = .27. Thus, with a training structure that included a mix of typical and atypical items and a young adult sample that showed a mix of prototype and exemplar strategy use, we found some evidence of both prototype and exemplar correlates in the VMPFC in young adults. In older adults, the overall pattern of findings was similar to young adults, with effects that were non-significantly weaker for both prototype and exemplar tracking.

Next, we tested for effects of older age and prototype vs. exemplar model in the anterior hippocampus. Additionally, we tested whether any effects in the anterior hippocampus differed significantly from those in the posterior hippocampus. Once again, there was no significant main effect of age, *F*(1,62) = .43, *p* = .52, *η_p_*^2^ =.01, nor significant main effects of model, *F*(1,62) = .82, *p* = .37, *η_p_*^2^ =.01, or hippocampal subregion, *F*(1,62) = .56, *p* = .46, *η_p_*^2^ =.01. None of the age interaction effects were significant (all F’s < .6, p’s > .46), and evidence for age differences in anterior hippocampus prototype correlates (BF_10_ = 0.3) and exemplar correlates (BF_10_ = 0.3) was weak. The age difference in anterior hippocampus prototype correlates remained non-significant even after accounting for age differences in motion via a multiple regression, *ß* = .08, *SE* = .09, *t* = 0.89, *p* = .38. Thus, although the prototype > baseline effect in the anterior hippocampus was significant only in the young adult sample, we did not find even moderate evidence for age differences in anterior hippocampus prototype correlates.

The model x hippocampal subregion interaction effect was marginally significant, *F*(1,62) = 3.75, *p* = .057, *η_p_*^2^ = .06. Although the interaction did not reach significance, the pattern was in the same direction as our prior work: prototype correlates (M = .08, SD = .23) were marginally stronger than the exemplar correlates (M = -.002, SD = .23) in the anterior hippocampus, *t*(63) = 1.79, *p* = .077, *d* = .22. In the posterior hippocampus, the exemplar correlates (M = .03, SD = .22) showed a minimal advantage over the prototype correlates (M = .02, SD = .22) that was not significant, *t*(63) = .14, *p* = .45, *d* = .02.

Taken together, our ROI results largely replicate what we have seen previously in young adults, with the exception of the VMPFC showing some signs of tracking exemplar information in addition to tracking prototype information. This may reflect the use of a training structure that contains less typical exemplars and promotes exemplar representations to a larger degree than our prior work. We newly show that older adults engaged the VMPFC to track prototype information while prototype correlates in the anterior hippocampus were not significant. Yet, the evidence in favor of age reductions in prototype-based activation in both regions was weak.

#### 3.2.3. Neural model fits: whole brain analyses

In addition to targeted tests of regions previously shown to track prototype information in young adults, we conducted exploratory whole-brain analyses to identify other regions that might support prototype-based and exemplar-based categorization. Figure 4 depicts regions tracking prototype and exemplar predictors across the entire sample with significant clusters identified in Table 3. Medial prefrontal cortex and the anterior cingulate along with temporal regions like middle temporal gyrus and temporal pole significantly tracked prototype predictors, which is consistent with prior work in young adults (Bowman and Zeithamova, 2018). However, there was also prototype tracking in several parietal regions like angular and supramarginal gyrus that we had not found previously. There was only one exemplar-tracking cluster that passed a whole-brain correction: in the VMPFC just posterior to the larger prototype-tracking cluster. Thus, it seems that there are mostly separate subregions of the VMPFC tracking prototype and exemplar information. When we examined prototype and exemplar tracking regions separately for each age group (Table 3), young adults generally showed activation across the regions that were significant across the entire sample for prototypes, but did not show suprathreshold activation for exemplars. For older adults, there were no clusters significantly tracking either the prototype or exemplar predictors. Thus, despite showing a similar pattern of activation as young adults in the ROI analysis, we do not see older adults having the same kind of robust prototype response across the brain as young adults. These weaker effects in older adults likely reflect the smaller sample size for the older group, the poorer fit of both the prototype and exemplar models we found from behavior, or some combination of these two factors.

**Figure 4.**
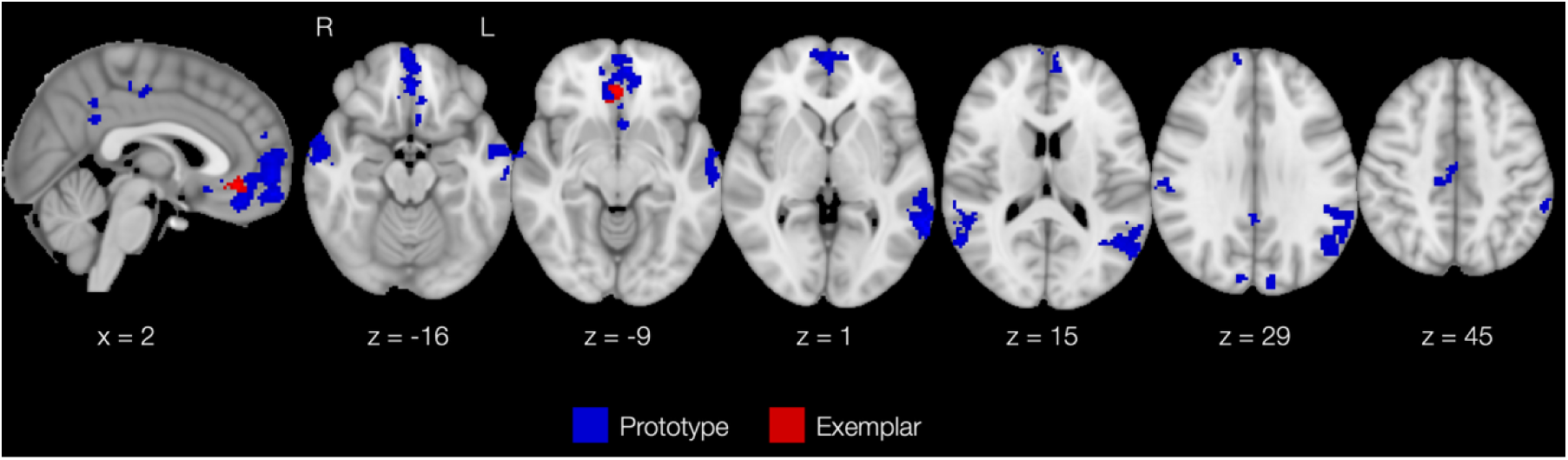
Whole-brain analysis depicting regions showing significant prototype > baseline (blue) and exemplar > baseline (red) across the entire sample. x-, y- and z-values are in MNI space. R = right, L = left.

**Table 3.**
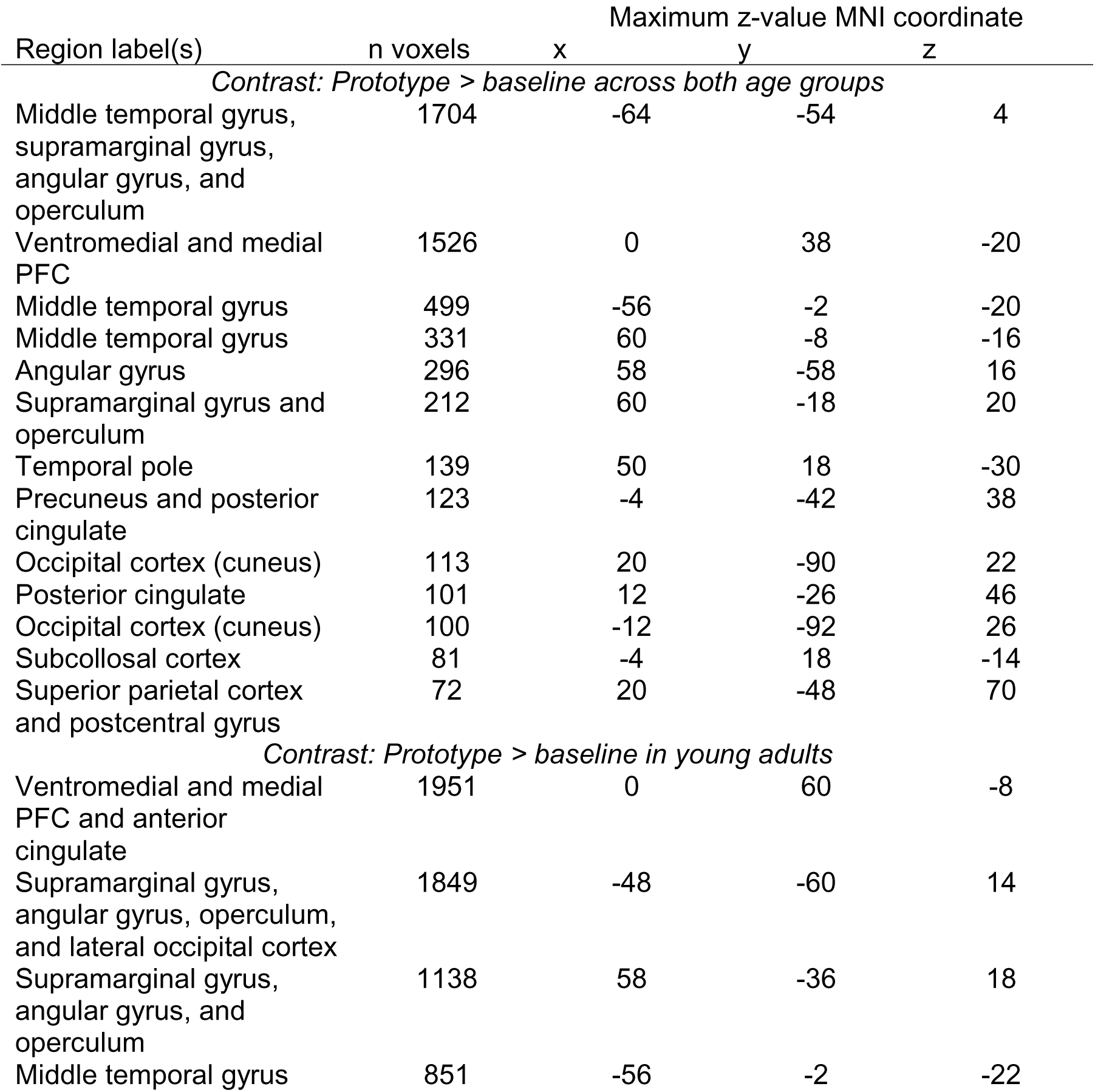
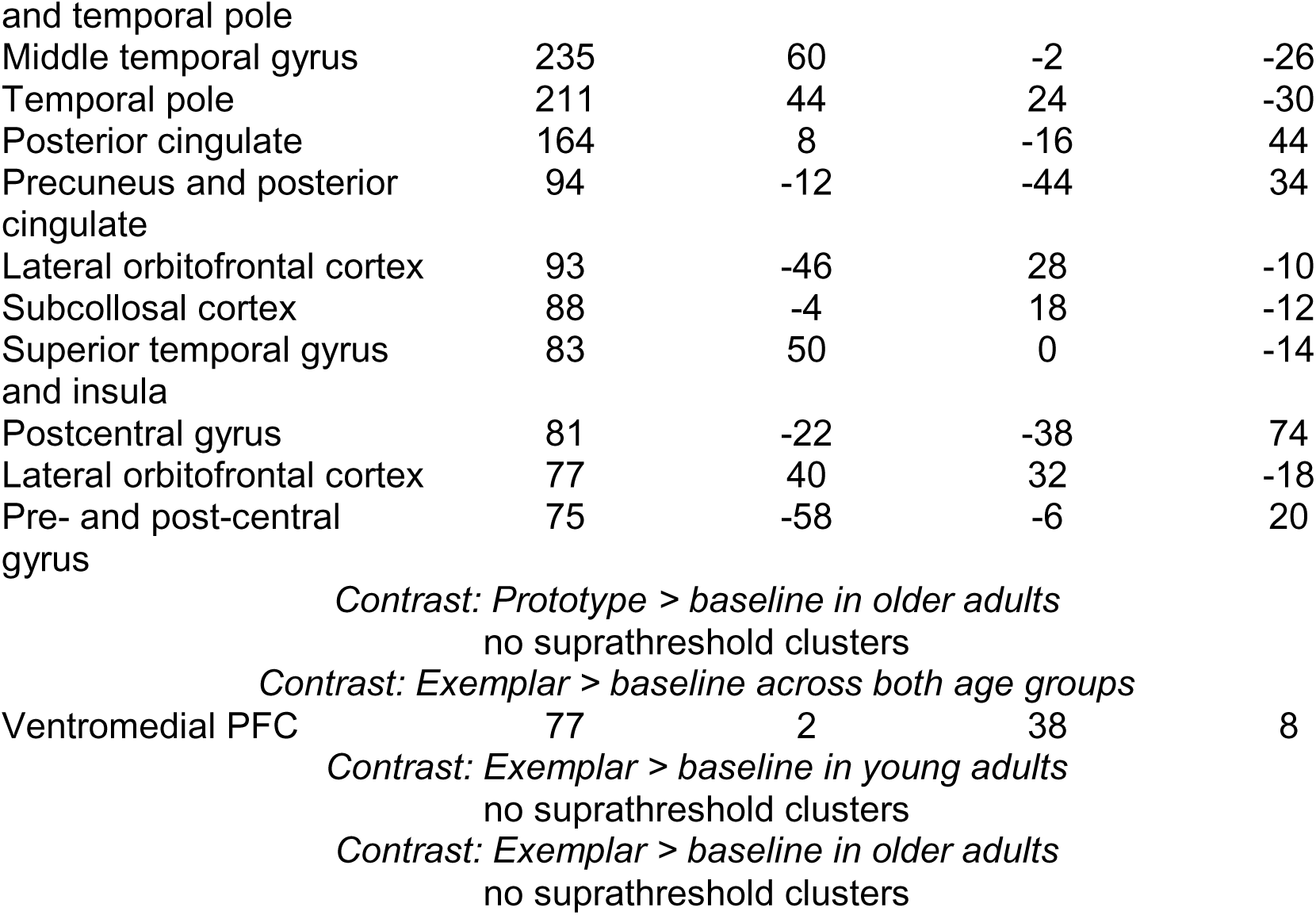
Cluster information for whole-brain prototype and exemplar correlates.

Next, we tested for significant age differences in prototype and exemplar activation across the whole brain (Table 4). There were no clusters that showed significant age differences in exemplar model correlates. Figure 5A depicts the single cluster in the right inferior frontal gyrus (IFG) showing stronger prototype-based activation in young adults compared to older adults. To test whether right inferior frontal gyrus significantly tracked prototype information in each age group and whether any prototype effects differed from exemplar tracking, we extracted the prototype and exemplar neural effects from each participant and computed one-sample t-tests comparing the prototype effect to baseline and paired samples t-tests comparing the prototype and exemplar effects in each age group (Bonferroni correction for four tests alpha = .0125). Note that we did not compute age comparisons at this stage since it was an age comparison that generated the cluster. The neural model fits are depicted in Figure 5B. The prototype tracking effect in young adults was positive but did not pass a correction for multiple comparisons, *t*(35) = 2.62, *p* = .013, *d* = .44, and the difference between prototype and exemplar correlates was not significant, *t*(35) = .437, *p* = .665, *d* = .07. There was significant prototype tracking in older adults, but in the negative direction, *t*(27) = -3.74, *p* < .001, *d* = -.71. That means that trials with higher levels of prototype similarity led to lower activation in this region. However, as with young adults, the difference between the prototype and exemplar correlates was not significant, *t*(27) = .588, *p* = .561, *d* = .11. Thus, there was an age difference in how the right IFG responded to the typicality of category members: stronger response to typical category members in young adults versus stronger response to atypical category members in older adults. Further, this effect was comparable regardless of the definition of category typicality: based on exemplar similarity or prototype similarity.

**Figure 5.**
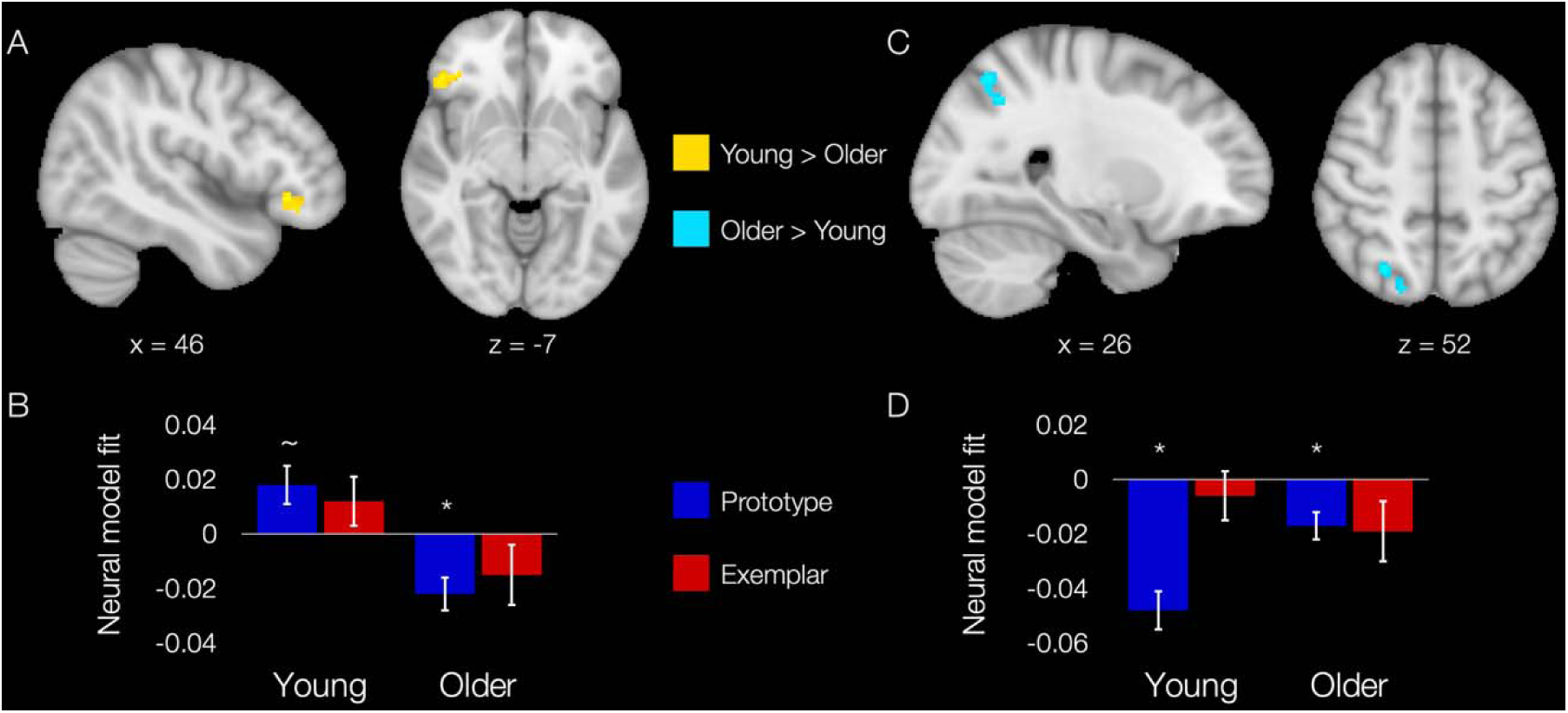
Whole-brain age differences in prototype correlates. A. A cluster (shown in two views) in the right inferior frontal gyrus showing young > older adult activity. B. Mean neural model fit (mean beta/variance) for the prototype and exemplar models in the right inferior frontal gyrus cluster. C. A cluster (shown in two views) in the right lateral occipital/superior parietal cortex showing older > young adult activity. B. Mean neural model fit (mean beta/variance) for the prototype and exemplar models in the right lateral occipital/superior parietal cortex cluster. For one-sample t-tests, ∼ represents a significant difference at alpha = .05, * at a corrected alpha of .0125.

**Table 4.**
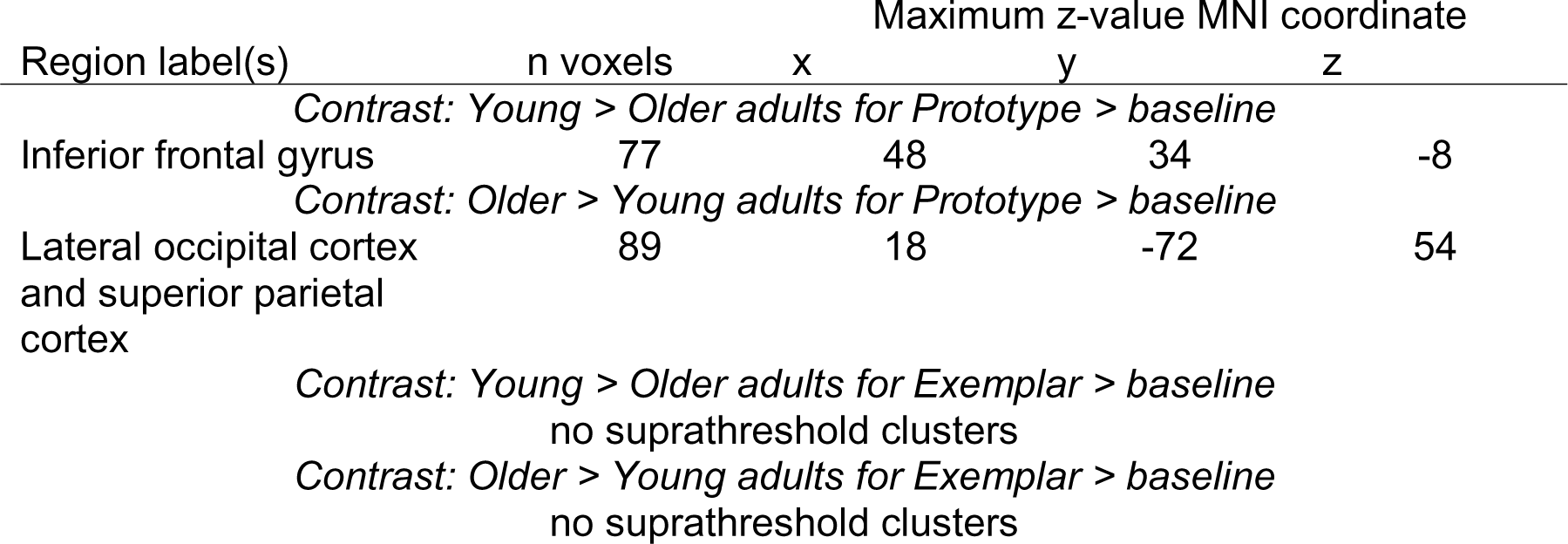
Cluster information for age differences in whole-brain prototype an exemplar correlates.

Figure 5C depicts the single cluster in the right lateral occipital/superior parietal cortex showing older adult > young adult activation for prototypes, and the corresponding neural model fits are depicted in Figure 5D. Yet, we see that the differences are in levels of negative prototype effects, meaning that activation was higher in this region for items further from the prototype (near category boundary). Both young and older adults showed significant negative prototype tracking in this region, young *t*(35) = -6.537, *p* < .001, *d* = -1.09, older *t*(27) = -3.213, *p* = .003, *d* = -.61. Young adults also showed significantly more negative tracking of prototype relative to exemplar similarity, *t*(35) = -2.795, *p* = .008, *d* = -.47, whereas older adults did not differentiate between prototype and exemplar tracking in this region, *t*(27) = .114, *p* = .910, *d* = .02. Thus, despite showing older > young adult activity, this occipitoparietal region was one that was again showing stronger prototype tracking in young adults, just in a negative direction.

### 3.3. Exploratory analysis of neural old vs. new differences

Some previous studies have identified exemplar correlates in the brain (Blank and Bayer, 2022; Bowman et al., 2020; Mack et al., 2013), but here we found a mix of prototype and exemplar strategies in behavior in young adults without strong corresponding exemplar representations in the brain. One key prediction of the exemplar model is that the individual training items should be stored in memory and should have a strong response during generalization since they are the basis of the exemplar representation. In line with this exemplar model prediction, we found a categorization advantage of old items over new items in behavior. We thus conducted an exploratory fMRI analysis testing old/new differentiation of items during the categorization test, which would suggest some degree of memory for the specific training items as predicted by the exemplar model. We further explored whether any old/new effects were stronger in young adults than in older adults, as would be expected from the mixture of prototype and exemplar strategies in this age group. To do so, we computed an exploratory univariate GLM where old and new stimuli and stimuli of different distances each had their own regressor (i.e., old distance 2 items, old distance 4 items, new distance 0 items (prototypes), new distance 1 items, etc.), leading to 7 separate regressors. All regressors had a duration of 5 seconds (as with the model-based analyses) and all onsets had a weight of 1 (no parametric modulation). We then contrasted old vs. new items at distance 2 and old vs. new at distance 4 to test for neural old/new discrimination while equating for distance from the prototype. We computed those contrasts within each age group and computed age difference (young vs. older) contrasts.

Table 5 presents the results of these old > new contrasts for young adults, older adults, and for age group comparisons. Group comparisons are also visualized in Figure 6. No old/new differences were found in older adults at either distance; we also did not find any reliable older > young effects. In contrast, young adults show a differentiation of old and new items for both distance 2 and 4, with several clusters showing reliable age differences in the old/new effects. At distance 2, the old > new contrast for distance 2 items yielded two suprathreshold clusters in young adults, and two cluster also reliably greater activation in young adults compared to older adults. Figure 6A depicts the two clusters showing suprathreshold young > older adult activation: one in left frontal pole and one that spanned the cuneus and precuneus. One-sample t-tests on the contrast weights averaged across all significant voxels showed a significant (Bonferroni correction for two tests alpha = .025) positive effect in young adults, *t*(35) = 2.86, *p* = .011, *d* = 0.45 and a significant negative effect in older adults, *t*(27) = -3.85, *p* = .007, *d* = -0.73. At distance 4, young adults showed significantly more activation in the old > new contrast in a set of occipital and parietal regions (Table 5). Further, activation across several overlapping occipital and parietal clusters showed greater old > new activity in young adults than in older adults (Table 5, Figure 6B). As with the contrast for distance 2 items, one-sample t-tests on the contrast weights averaged across all significant voxels showed a significant (Bonferroni correction for two tests alpha = .025) positive effect in young adults, *t*(35) = 4.15, *p* < .001, *d* = 0.69 and a significant negative effect in older adults, *t*(27) = -4.47, *p* < .001, *d* = -0.85. Thus, unexpectedly, both young and older adults showed neural discrimination of old versus new items, but in different directions. Across primarily occipital and parietal regions, young adults showed enhanced processing of the old training items, whereas older adults showed a novelty effect.

**Figure 6.**
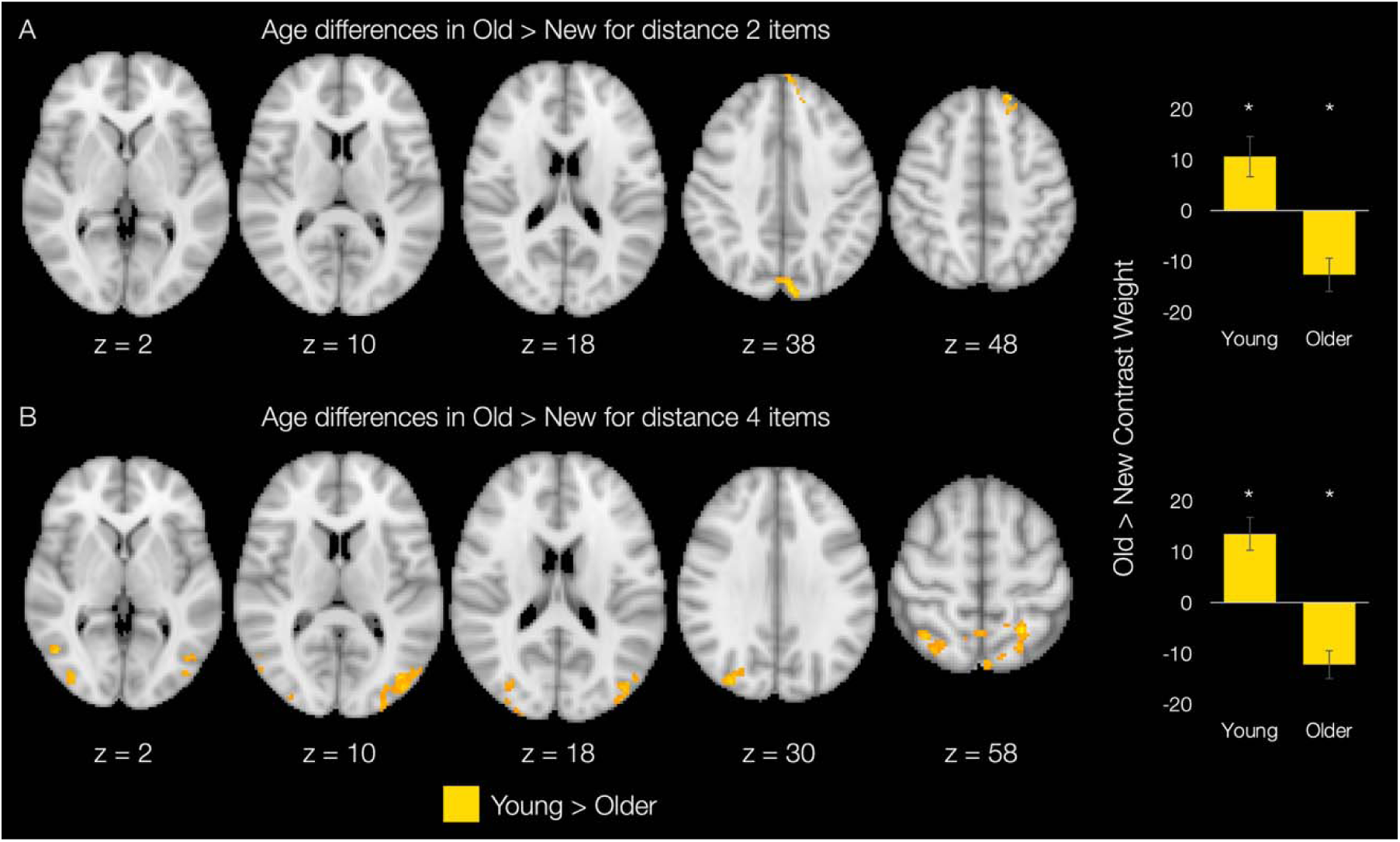
Whole-brain age differences in old vs. new discrimination. A. Distance 2 items (category-typical items close to the category prototypes): two clusters showed greater old vs. new discrimination in young compared to older adults. B. Distance 4 items: (less typical items near category boundary): nine clusters across occipital and parietal cortex showed greater old vs. new discrimination in young compared to older adults. No suprathreshold clusters showed the opposite age difference (older > young) for either distance 2 or distance 4 items. Contrast weights are averaged across all voxels showing significant young > older activation in a given contrast.

**Table 5.** Cluster information for whole-brain old > new analysis for distance 4 items.

| Region label(s) | n voxels | Maximum z-value MNI coordinate |  |  |
| --- | --- | --- | --- | --- |
|  |  | x | y | z |
| *Contrast: Old > New – distance 2 items in young adults* |  |  |  |  |
| Pre/post-central gyrus | 85 | -58 | -14 | 42 |
| Frontal pole | 67 | -44 | 44 | -12 |
| *Contrast: Old > New – distance 2 items in older adults* |  |  |  |  |
| no suprathreshold clusters |  |  |  |  |
| *Contrast: Young > Older adults for Old > New – distance 2 items* |  |  |  |  |
| Frontal pole | 84 | -18 | 44 | 48 |
| Cuneus and precuneus | 68 | -6 | -80 | 36 |
| *Contrast: Older > Young adults for Old > New – distance 2 items* |  |  |  |  |
| no suprathreshold clusters |  |  |  |  |
| *Contrast: Old > New – distance 4 items in young adults* |  |  |  |  |
| Lateral occipital cortex | 155 | 30 | -68 | 8 |
| Superior parietal cortex | 153 | 34 | -54 | 62 |
| Superior parietal cortex | 98 | -26 | -54 | 66 |
| Precuneus | 91 | 2 | -52 | 66 |
| Precentral gyrus | 89 | 30 | -20 | 70 |
| Lateral occipital cortex | 75 | -40 | -72 | 10 |
| Precuneus | 60 | 2 | -46 | 50 |
| Lateral occipital cortex | 56 | 52 | -74 | 10 |
| *Contrast: Old > New – distance 4 items in older adults* |  |  |  |  |
| no suprathreshold clusters |  |  |  |  |
| *Contrast: Young > Older adults for Old > New – distance 4 items* |  |  |  |  |
| Lateral occipital cortex | 331 | -40 | -78 | 12 |
| Superior parietal cortex | 320 | -22 | -54 | 56 |
| Precentral gyrus | 208 | 16 | -18 | 74 |
| Precuneus | 161 | -2 | -52 | 64 |
| Lateral occipital cortex | 148 | 36 | -82 | 28 |
| Lateral occipital cortex | 82 | 38 | -86 | 4 |
| Superior parietal cortex | 76 | 36 | -54 | 62 |
| Precuneus | 56 | -4 | -74 | 56 |
| *Contrast: Older > Young adults for Old > New – distance 4 items* |  |  |  |  |
| no suprathreshold clusters |  |  |  |  |

## 4. Discussion

In the present study, we tested whether the VMPFC and anterior hippocampus support prototype representations in older adults as they do in young adults. As expected, older adults were more likely than young adults to be best fit by the prototype model, although the overall fit of both models was poorer in older compared to young adults. In the brain, we found that both young and older adults showed significant prototype tracking in the VMPFC, but the prototype effect in the anterior hippocampus was only significant in young adults. Across both regions, evidence for age differences in prototype correlates was weak, but did emerge in separate inferior frontal and occipitoparietal regions. The relatively preserved VMPFC prototype processing in older adults was coupled with minimal age deficits in behavioral generalization performance, demonstrating that prototype representations can support successful generalization in older adults. We also found age differences in neural discrimination of old and new items in occipital and parietal regions, with young adults showing increased activation for old items and older adults showing increased activation for novel items. Thus, concept generalization in older adults is supported by an increased reliance on prototype representations, engagement of the VMPFC to track prototype information, and a potential shift towards novelty processing in occipitoparietal regions.

As in our prior behavioral study that used the same category structure and a similar task procedure (Bowman et al., 2022), we found that older adults’ categorization responses tended to be best fit by the prototype model, and that older adults were more likely to be best fit by the prototype model than young adults were. We also replicated two other behavioral patterns: 1) there was a smaller age deficit in learning typical training items that were close to category prototypes (10% accuracy difference) compared to more atypical items near the category boundary (20% accuracy difference), and 2) age deficits in category generalization were minimal despite robust age deficits in category training. Together, these findings suggest that older adults’ memory is sufficiently equipped to extract the commonalities among members of the same category, which is adequate for learning typical category members and for generalization. In contrast, memorization of specific training items—a strategy that young adults seem to use to boost training performance for atypical items—may be hindered in aging. These findings are also consistent with memory and aging findings outside of category learning, which have shown that the ability to remember the general meaning or ‘gist’ of an experience is intact in older age (Brainerd and Reyna, 2015, 2002) while memory for specific, unique details of individual experiences declines (Bowman et al., 2019; Bowman and Dennis, 2015; Gallo et al., 2006; Koutstaal and Schacter, 1997). Thus, across multiple memory domains, the older brain seems to prioritize building abstract, generalizable knowledge over remembering individual experiences.

While we have previously shown prototype correlates in the anterior hippocampus in young adults, there was only a non-significant trend toward prototype representations in older adults. Yet, the age difference was not significant and evidence in favor of an age difference was quite weak. Thus, the role of the anterior hippocampus in prototype-based categorization in older age is somewhat equivocal. The hippocampus has long been known to support rapid learning and retrieval of unique experiences (Chadwick et al., 2010; Fortin et al., 2004; Leutgeb et al., 2007; Scoville and Milner, 1957), but more recent work has also identified a role for the hippocampus in integration of related experiences and in generalization (Bowman et al., 2020; Bowman and Zeithamova, 2018; Schlichting et al., 2015; Shohamy and Wagner, 2008; Zeithamova et al., 2012). It is thought that the hippocampus can form relationships between separate but related experiences very rapidly before they have had time to be consolidated into neocortical networks (Schapiro et al., 2017). Here, we find that, although the hippocampus is known to undergo structural and functional decline in older age (Driscoll et al., 2003; Leal and Yassa, 2015; Lister and Barnes, 2009), strong evidence for age differences in its contributions to prototype-based generalization did not emerge. One possibility is that hippocampal functions like pattern separation (Yassa et al., 2011) and pattern completion (Wynn et al., 2021) are both differentially affected by aging and differentially contribute to prototype-based generalization, leading to the weak level of age-related hippocampal processing decline found here. Future work directly comparing hippocampal prototype processing with age differences in pattern separation and completion would be helpful in further elucidating the role of the hippocampus in generalization in older age.

Further, both young and older adults showed significant prototype tracking in the VMPFC, which is consistent with prior work in young adults alone (Bowman et al., 2020; Bowman and Zeithamova, 2018; Liu et al., 2025; Blank and Bayer, 2022). In addition to tracking prototype information, the VMPFC is known to integrate across related experiences (Audrain and McAndrews, 2022; Schlichting et al., 2015; Spalding et al., 2018; Tompary and Davachi, 2017; Zeithamova et al., 2012) and support schema-based memory (Ghosh et al., 2014; Reagh and Ranganath, 2023; van Kesteren et al., 2010). The VMPFC may therefore broadly support formation of abstract representations that summarize across experiences and apply those abstract representations to new situations.

While lateral prefrontal cortex tends to show age-related differences in memory function (Davis et al., 2008; Kim and Giovanello, 2011; Rajah and D’Esposito, 2005), work on potential age differences in medial prefrontal function have been more mixed (Gutchess, 2014; Jobson et al., 2021). Here, we show that prototype-based generalization in the medial prefrontal cortex did not differ significantly between young and older adults. Our results thus align with other work showing comparable medial PFC function (Leshikar and Duarte, 2014), which extend to other domains like social processing (Cassidy et al., 2012; Gutchess et al., 2007). Thus, older adults’ ability to maintain memory performance in terms of category generalization, schema-based memory, and gist-based memory may be due in part to relatively intact medial PFC function.

We also found novel evidence of *exemplar* correlates in the VMPFC in addition to the prototype correlates we have seen previously. There was a trend toward significant VMPFC exemplar tracking in young adults when we considered the VMPFC as a whole, and there was also a cluster in the VMPFC that was significant in the whole-brain analysis across age groups. The exemplar-tracking cluster was smaller than the prototype-tracking cluster we also identified at the whole-brain level, and the exemplar and prototype clusters overlapped only minimally. While the exemplar effect was not significant in older adults alone in the ROI analysis, there was also no age difference in the VMPFC exemplar effect. Thus, it seems that the VMPFC is capable of maintaining both abstract and item-specific representations during categorization and does so in a similar manner across age groups. This finding is also consistent with findings showing that the VMPFC can represent categories at multiple levels of complexity (Mack et al., 2020), that it can contain a mix of integrated and separated representations in an associative inference task (Schlichting et al., 2015), and with its role as a hub region that integrates many sources of information (Buckner et al., 2009). Further, work on schema memory shows that the VMPFC is involved in tracking the congruity of individual items with a broader context, suggesting role of the VMPFC both in representing the larger, abstract context and in integration of individual items into that context (van Kesteren et al., 2012). Yet, we have also seen prototype correlates most strongly and most consistently in this region (Bowman et al., 2020; Bowman and Zeithamova, 2018; Liu et al., 2025), suggesting that its primary role is in representing the abstract, generalized information, at least in the context of categorization tasks.

We did not find significant age differences in prototype correlates in the hippocampus or VMPFC, but we found two regions showing age differences in our whole-brain analysis. One region in the prefrontal cortex seemed to track item typicality generally, irrespective of whether category typicality was based on prototype- or exemplar-similarity. Interestingly, the age difference in this region was driven by a flip of the direction of the effect: positive tracking in young adults and negative tracking in older adults. In other words, young adults were activating this region more strongly for items near category center while older adults were activating this region more strongly for atypical items near category boundary. Thus, there seems to be an age differences in whether prefrontal cortex prioritizes more typical versus atypical feature combinations when making categorization judgments.

A region in right occipitoparietal cortex showed stronger negative prototype tracking in young adults than in older adults: this region showed higher activation for less prototypical category members (items near category boundary) and that effect was stronger in young than older adults. This occipitoparietal region also differentiated prototype from exemplar correlates in young but not older adults. Thus, there seem to be age differences in visual processing of category members, with young adults showing stronger responses to less prototypical category members, potentially reflecting visual attention to additional visual features when the most highly attended features are not sufficient to determine category membership (Nosofsky, 1986). Another possibility is that this activation functions as a novelty signal for less typical items because their combination of features is less familiar (Kafkas and Montaldi, 2018). That older adults showed reduced tracking of prototype information in this visual region suggests there may be age differences in the level of visual detail in the prototypes, which can be explored in future studies.

Lastly, although we found that older adults were more likely than young adults to be best fit by the prototype model, the fit of the prototype model was poorer in older adults, and we did not see age-related increases in prototype correlates in the brain. This pattern indicates that the age-related increase in prototype reliance could be driven by a lack of exemplar processing rather than an increase in prototype processing. A deficit in exemplar processing could leave prototypes as the only viable option for representing categories, whereas young adults may have the flexibility to use a mix of representations. We saw some evidence for this proposal in that young adults were more likely than older adults to be best fit by the exemplar model in behavior. Yet, we did not find widespread exemplar model correlates in the brain in either age group, and using neural discrimination of old versus new items as an alternative index of exemplar processing showed a shift toward novelty processing in older adults rather than a lack of old/new discrimination. Future research will be necessary to test the extent to which this shift toward novelty processing in the brain contributes to exemplar- versus prototype-based categorization, potentially linking individual differences in neural old/new discrimination to age differences in the degree of prototype advantage in behavior with a sample powered for such analyses.

While follow-up studies will be needed to further establish concept-learning mechanisms in older adults, our findings have several implications that could be useful for supporting concept learning in older adults. Along with our prior work (Bowman et al., 2022, 2021), we show that identifying appropriate training regimes for older adults is likely to be the largest hurdle, whereas concept generalization will likely require less intervention and support. The lack of age differences in learning typical training examples points to the possibility that training regimes consisting of typical category members will be the most fruitful for promoting concept learning in older adults. Finally, to the extent that prototype learning systems show fewer age differences than systems for individuating category members, interventions may be more effective when they leverage this prototype system and focus on promoting abstract knowledge. Following these principles has the potential to help older adults learn new concepts as they enter the world and engage in new activities that may enrich their lives.

## 5. Conclusion

We tested for age differences in the neural basis of prototype-based category generalization and found that young and older adults recruited the VMPFC to a comparable degree to support prototype representations Thus, older adults are able to engage the VMPFC to support abstract concept representations. This relatively preserved VMPFC function was coupled with a lack of age deficits in generalization performance. These findings raise the possibility of leveraging prototype-based processing to support novel concept learning among older adults.

## 6. Acknowledgements

This work was supported by the National Institutes of Health through grants R01-NS112366 (Zeithamova) and F32-AG054204 (Bowman) and by the Lewis Family Endowment, which supports the Robert and Beverly Lewis Center for Neuroimaging at the University of Oregon.

## Appendix A

### Prototype and exemplar model fitting to behavioral responses

The prototype and exemplar models were fit to trial-by-trial categorization data in individual participants. The prototype model assumes that items are categorized based on their similarity to category prototypes. Prototype similarity is computed as an exponential decay function of the distance in psychological space between the to-be-categorized item and the prototype of a given category (Shepard, 1957). Estimated model parameters include the amount of attention paid to individual features (attention weights) and the steepness of the function relating distance to subjective similarity (sensitivity). Formally, the similarity of item x to the category A representation is computed as:

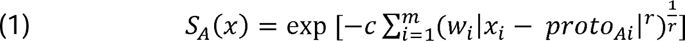

where S_A_(x) is the similarity of item x to the category A representation, x_i_ is the value of the i-th dimension of item x, m is the number of feature dimensions (m = 10 in the present study), proto_A_ is the prototype of category A, r is the distance metric (fixed at 1 for the city-block metric in the present study), w is a vector with attention weights for each of the 10 stimulus features (constrained to sum to 1), and c is the sensitivity (constrained to be between 0-100).

The exemplar model is the same except that the underlying similarity computation is the summed similarity across training exemplars from a given category, but it includes the same estimated attention weights and sensitivity parameters. Similarity of item x to the category A representation is computed as:

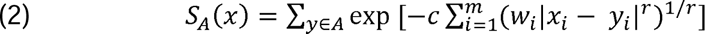

where y represents the individual training stimuli from category A, and the remaining notations are as in Equation 1.

For both models, the probability of assigning a given stimulus to category A is equal to the similarity to category A divided by the summed similarity across categories A and B:

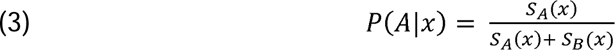

For each trial and each model, we estimated the probability of the participant having made their observed response. We then generated an error metric, computing the negative of log-transformed response probabilities (negative log likelihood) across all trials in the categorization phase and summing them. We used standard maximum likelihood methods implemented with fminsearch in MATLAB (MathWorks) to minimize this error metric separately for each model in each participant, adjusting the attention weights and sensitivity values to minimize model error. After minimization, the resulting negative log likelihoods were carried forward for model selection and the denominator from Equation 3 (summed similarity) for each model was used as a parametric modulator in fMRI analyses.

### Model selection

Our model selection approach had two steps: we first determined whether each model (prototype, exemplar) fit better than a model assuming random responding, and we then determined whether either the prototype or exemplar model fit better than the other. For both steps, we used a permutation approach that we have used previously (Bowman et al., 2022, 2020; Bowman and Zeithamova, 2023, 2020, 2018, in press) and described in detail (Zeithamova et al., 2025).

To determine whether each model fit better than one would expect by chance, we randomly shuffled each participant’s responses, maintaining any overall response bias but removing any relationship between the stimuli and the responses. We then fit each model to the permuted data and stored the resulting negative log likelihood value. We repeated this process 10,000 times, generating a distribution of negative log likelihood values that one would expect from chance responding. We then compared the fit of each model to the observed responses. Each model was considered to fit better than chance if the observed negative log likelihood value was smaller than 95% or more of the fits in the null distribution (i.e., alpha = .05, one-tailed). In the present study, both the prototype and exemplar model fit better than one would expect by chance for all included participants.

To determine whether the prototype model fit better than the exemplar model or vice versa, we computed a null distribution of differences in model fits using the fits to permuted responses from above. We subtracted the null prototype model fits from the null exemplar model fits and divided by their sum (i.e., exemplar fit – prototype fit / exemplar fit + prototype fit). Dividing by the sum of model fits accounts for how well the models fit overall in determining whether the difference in fits is significant. We computed the same difference for the fit of the prototype and exemplar models to the observed data. We then tested whether the difference in prototype and exemplar model fits was greater than would be expected by chance based on the null distribution of model fits (alpha = .05, two-tailed). Participants for whom the prototype model fit significantly better than the exemplar model were given the ‘prototype’ strategy label. Participants for whom the exemplar model fit significantly better than the prototype model were given the ‘exemplar’ strategy label. Participants for whom neither model significantly outperformed the other were given the ‘similar’ fits strategy label, indicating that we did not have enough evidence to say that one model fit better than the other.

